# How does the brain represent the semantic content of an image?

**DOI:** 10.1101/2021.06.23.449677

**Authors:** Huawei Xu, Ming Liu, Delong Zhang

**Affiliations:** Key Laboratory of Brain, Cognition and Education Sciences, Ministry of Education, Guangzhou 510631, China; School of Psychology, Center for Studies of Psychological Application, and Guangdong Key Laboratory of Mental Health and Cognitive Science, South China Normal University, Guangzhou 510631, China; Plateau Brain Science Research Center, South China Normal University/Tibet University, Guangzhou/Lhasa 510631/850000, China

**Keywords:** Deep neural networks, Neural style transfer, Early visual areas, Ground cognition, Encoding models, Representational similarity analysis

## Abstract

Using deep neural networks (DNNs) as models to explore the biological brain is controversial, which is mainly due to the impenetrability of DNNs. Inspired by neural style transfer, we circumvented this problem by using deep features that were given a clear meaning—the representation of the semantic content of an image. Using encoding models and the representational similarity analysis, we quantitatively showed that the deep features which represented the semantic content of an image mainly predicted the activity of voxels in the early visual areas (V1, V2, and V3) and these features were essentially depictive but also propositional. This result is in line with the core viewpoint of the grounded cognition to some extent, which suggested that the representation of information in our brain is essentially depictive and can implement symbolic functions naturally.

## 1. Introduction

Deep neural networks (DNNs) for image recognition provided an important tool for understanding the nature of visual object recognition [1–5]. This is not only because DNNs are currently the only models known to achieve near 5 human-level performance in object recognition, but also because they have the properties such as the hierarchical organization and the parallel distributed processing which are similar to the visual ventral stream—key circuits that underlie visual object recognition [6, 7]. Using DNNs as computational models, researchers found that DNNs could predict brain activity of visual processing across multiple hierarchical levels at unprecedented accuracy for both macaque [8–11] and human [12–15] wherein later layers in DNNs better predict higher areas of the visual ventral stream. The predictive power of DNNs made “mind-reading” possible [16–18] and promoted the integration of neuroscience and artificial intelligence [19, 20].

Besides the predictive power, an ideal model should also possess the explanatory power, which means that we should understand how the model works [21]. This is not the case of DNNs. DNNs are essentially black boxes and we can not understand how the input data were transformed into model output for now[22]. This is mainly due to the end-to-end learning and the huge number of parameters in DNNs (the complex architectures of DNNs). For example, AlexNet has about 60 million self-learned parameters [23] and VGG16 has 138 million self-learned parameters [24]. Even though we know the exact value of all parameters for each input, we still can not understand what do these parameters really mean. So using DNNs as models to explore the biological brain is something like replacing a black box with another, the lack of explanatory made it controversial [1]. To open the black box and look inside, researchers developed methods such as network dissection [25] and visualization [26–30], and experimented with network architecture [31], learning algorithm [32, 33], and input statistics [34]. But none of them can directly explain the meaning of the parameters (deep features) learned by DNNs.

However, an interesting and successful application of DNNs may give us a hint about the meanings of some deep features. Neural style transfer (NST) is a computer vision technique that allows us to render the semantic content of an image in the style of another [35–37]. Using NST, for example, we can blend a photo with van Gogh’s “Sunflowers” to get a new image which preserve the content of the photo but looks like if it was painted by van Gogh. According to the seminal work of [36], the implementation of the original NST algorithm was based on a DNN optimized for object recognition—VGG19. This process took two images, a content image and a style image. First, two images were fed into the pre-trained VGG19 model to extract feature maps, respectively. Second, the feature maps of the conv4_2 layer of the content image were selected as the semantic content representation. Third, the feature maps of the conv1_1 layer, conv2_1 layer, conv3_1 layer, conv4_1 layer, and conv5_1 layer of the style image were selected to compute the Gram matrix as the style representation. Last, through jointly minimizing the distance of the feature representations of a white noise image from the content representation and the style representation (feature inversion using the same VGG19 model), a new image was generated which simultaneously match the content of the content image and the style of the style image. The key to NST lies in the ability to extract representation from an image which explicitly separate image content from style [36].

In this study, we focused on the deep features extracted from the conv4_2 layer of the VGG19, which were selected as the representation of the semantic content of an image in the original NST algorithm. Under the framework of voxel-wise encoding models [38–40], it gave us an opportunity to explore the question of how does the brain represent the semantic content of an image. Although the original NST algorithm was effective which led to lots of follow-up studies and many successful industrial applications (e.g., Prisma), it was still hard to define what is the semantic content of an image and what is the style of an image—“The separation of image content from style is not necessarily a well defined problem. … In our work we consider style transfer to be successful if the generated image ‘looks like’ the style image but shows the objects and scenery of the content image. We are fully aware though that this evaluation criterion is neither mathematically precise nor universally agreed upon” [36]. For this question, we used representational similarity analysis [RSA, 41–43] to explore the representational similarity between the representation of the semantic content of an image and other representations such as the representation of semantics and the representation of Gabor features. So we explored two questions in this study: how does the brain represent the semantic content of an image and what is the semantic content of an image(Fig.1).

**Figure 1:**
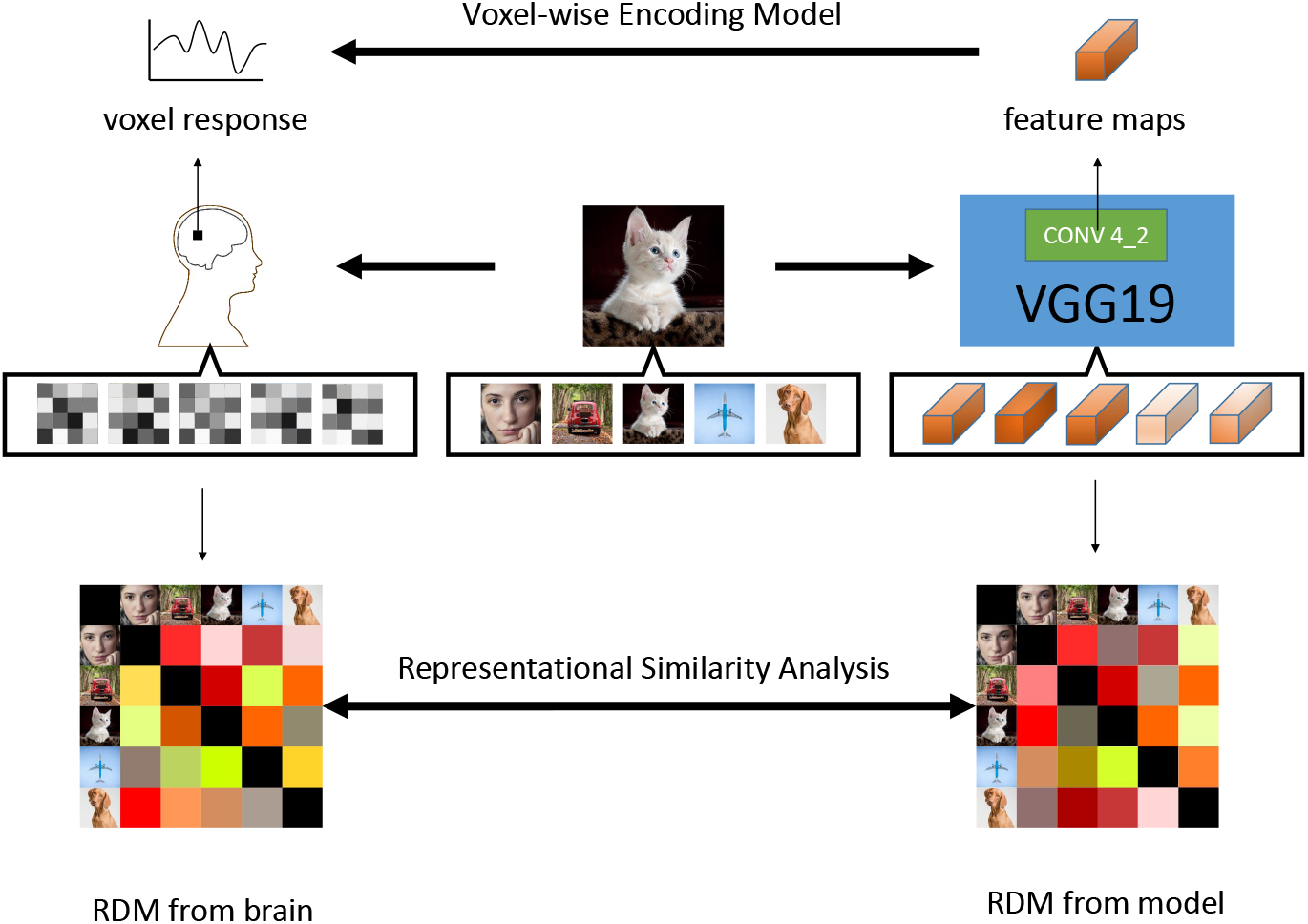
The Experimental Workfolw: encoding models and representational similarity analysis.

## 2. Materials and methods

### 2.1. Data

We used the fmri data that was originally published in [16], which can be downloaded from https://github.com/KamitaniLab/GenericObjectDecoding. The data was obtained from two fMRI experiments for each of 5 subjects: an image presentation experiment and an imagery experiment. There were two sessions in the image presentation experiment—the training session and the testing session. In the training session, subjects viewed 1200 images from 150 categories (8 images from each category) as each image presented once. In the testing session, subjects viewed 50 images from 50 categories (one image from each category) as each image presented 35 times. In the imagery experiment, subjects were asked to imagine about 50 nouns from 50 categories (one noun from each category) as each noun presented 10 times. The categories used in the imagery experiment were the same as those in the testing session, which were not used in the training session. All the images (1250 natural images with the resolution of 512 × 512 × 3) and the corresponding categories (200) were collected from ImageNet [44]. In addition, there were standard retinotopic mapping experiment and functional localizer experiment to identify lower visual areas (V1, V2, V3, and V4) and higher visual areas (LOC, FFA and PPA) for each participant. The details of the experimental design, MRI acquisition protocol, and preprocessing of the fMRI data could be found in [16].

Before further analysis, we averaged the repeated trials in the testing session and the imagery experiment. First, we standardized the fMRI data from the training session. The mean and standard deviation of the training set were then used to standardize the testing set (from the testing session) and the imagery set (from the imagery experiment). After that, we performed trial-averaging for the testing set and the imagery set to improve the signal-noise ratio. Because of trial-averaging, there were statistical difference between the training set and the other two. So we rescaled the averaged data by a factor of 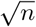 where n is the number of trials averaged [17].

### 2.2. Encoding Models Based On Deep Features

To probe how does the brain represent the semantic content of an image, we used voxel-wise encoding models. Such models were constructed separately for each voxel in each individual, so our analysis was also individual-based. There were two steps to construct voxel-wise encoding models [40]: the first step was a nonlinear transformation from a stimulus space to a feature space; the second step was a linear transformation from the feature space to a voxel space.

As the first step, we got deep features which represented the semantic content of an image. We employed the pre-trained VGG19 model based on the open source machine learning framework of PyTorch [45] to extract the feature maps of the conv4_2 layer as the image content representation. After image preprocessing (such as image scaling and cropping, more details can be found from https://pytorch.org/hub/pytorch_vision_vgg/), images from the training data session and the testing data session were fed into the VGG19 model and the feature maps of the conv4_2 layer were extracted, respectively. As a result, the size of the training feature maps was [1200, 512, 28, 28] ([the number of images, the number of kernels, the height of feature map, the width of feature map]), and the size of the testing feature maps was [50, 512, 28, 28].

As the second step, we constructed linear regression models to predict the brain activity evoked by an image from the features which represented the semantic content of the same image. For each voxel, the model can be expressed by

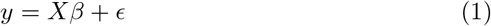

Where *y* is a measured voxel response and *X* is features extracted from image which elicited the response of the voxel.

Just as each neuron has its own receptive field, each voxel has its own population receptive field [46]. It means that a voxel only responds to the features in its population receptive field. So there was no need to put all the features into the model (The number of features for each image is 401408, It will pose a problem known as the curse of dimensionality if we put all the features into the model). And the features we extracted from the VGG19 model were naturally organized into feature maps that preserved the topology of stimuli. For example, the size of the feature maps for each image was 512 × 28 × 28. It could be seen as 784 (28 × 28) spatial locations with 512 features at each spatial location. The spatial arrangement of 784 locations (28 × 28) preserved the topology of the original image (512 × 512). According to the model of the population receptive field, receptive fields are center-surround organized and features at the center of a receptive field make the largest contribution to the activity measured in the voxel [47]. So we only used 512 features at one of the 784 locations as *X* for each voxel (To simplify the model and reduce computation time, we ignored the surround of receptive fields and assumed that each location is a candidate for the center of a receptive field). To find the best center for each voxel, we constructed separate linear regression models for each location/voxel combination.

We used the training data to fit models. For each model, *X* was the features from one of the 784 locations represented by a 1200 × 512 matrix (1200 images), *y* was a measured voxel response represented by a 1200 × 1 matrix. It was reasonable to assume that a voxel only responded to a fraction of the *X*. So we estimated regression coefficients of each model using lasso regression:

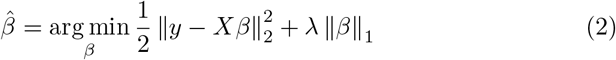

where lambda is a complexity parameter that controls the amount of regularization. To accelerate model fitting, we used the function “lasso_gpu” from a MATLAB package developed by [48], which can be efficiently implemented in parallel on a GPU. The optimal value of lambda was selected from a lambda sequence with 10 lambdas: 2^−1^, 2^−2^, 2^−3^, ⋯, 2^−10^ (the first lambda was set using the function “calculate_lambda_start” from the same MATLAB package and the lambda range was set to guarantee that the last lambda is not chosen as the best one). We chose the best lambda and the best location for each voxel using five-fold cross-validation with the coefficient of determination (*R*^2^).

Once fitted, encoding models were evaluated using the testing sets and the imagery sets, respectively. For each voxel, we defined the model’s prediction accuracy as the Pearson’s correlation coefficient (*r*) between the measured voxel response and the response predicted by the model. The significance of the correlation was assessed by a permutation test with 10000 permutations (*p* < 0.01) and FDR was applied for multiple comparison correction [49]. For each ROI, we used the number of survived voxels within the ROI and the meidan r as measurements for prediction accuracy.

Another measurement for prediction accuracy was decoding performance—identifying stimuli from measured brain activity [50]. First, we used survived models to predict the voxel activity pattern from the test feature maps for each of the 50 stimuli. Second, we calculated the Pearson’s correlation coefficient between the predicted voxel activity pattern and the measured voxel activity pattern for each stimulus/stimulus combination. The stimulus whose predicted voxel activity pattern was most correlated with the measured voxel activity pattern of itself was regarded as correct decoding. We defined the identification accuracy as the percentage of stimuli that are correctly identified from the testing data or the imagery data.

After model evaluation, we explored the relationship between features and voxels through regression coefficients of survived models. Because of the lasso regression, some of regression coefficients in each model were set to zero automatically. The nonzero coefficients indicated that which features were related to the activity of a voxel and how are they related. For each subject, we described the relationship between features and voxels from two complementary perspective—the perspective of features and the perspective of voxels. From the perspective of features, we used the number of voxels each feature related and the location distribution of these voxels in different ROIs to analyse the property of different features. From the perspective of voxels, we used the number of features each voxel related and the location of the voxel to analyse the property of different ROIs.

Furthermore, the survived models also provided information about the population receptive fields of voxels. For each model, the *X* was selected from one of the 784 locations (28 × 28). The location of the *X* could be seen as the center of the population receptive field of the corresponding voxel on the feature maps. We used heatmap to visualize and explore the distribution of the receptive fields of survived voxels for each subject.

### 2.3. Encoding Models Based On Gabor Features

To compare with encoding models using the deep features, we also constructed encoding models based on Gabor features. We got gabor features of stimuli according to the method of [51]: Firstly, a Gabor Wavelet Pyramid [GWP, 52] model was used to get original Gabor features from stimuli (six spatial frequencies: 1, 2, 4, 8, 16, and 32 cycles/FOV; eight orientations: 0°, 22.5°, 45°, …, and 157.5°; and two phases: 0° and 90°; The FOV covered full of a image and all images were downsampled to 128 × 128 pixels); Secondly, the absolute values of the projections of each quadrature wavelet pair were averaged to get the contrast energies; Thirdly, the contrast energies were normalized to linearize the relationship between contrast energies and voxel responses (each contrast energy was divided by the sum of the contrast energy and the median of all contrast energies in the training set which were at the same position and the same orientation); Fourthly, the normalized contrast energies at the same position (eight orientations) were averaged to reduce the dimension of features; Lastly, the average luminance of stimuli were also added into Gabor features. As a result, each stimulus had 1366 features. After that we used the same method as described above to construct linear regression models from Gabor features to voxel responses.

### 2.4. Representational Similarity Analysis

To explore the question of what is the semantic content of an image, we used RSA, which characterized the representational geometry of a brain region or computational model (using a set of stimuli) by representational distance matrix (RDM) and compared RDMs to explore the representational similarity between different brain regions or brain regions and computation models. Three types of RDMs (the candidate RDMs) were constructed to compare with the RDM of the layer conv4_2 (the reference RDM). In accordance with the analysis using encoding models, the RSA was also based on individuals.

The first type were RDMs derived from the pre-trained VGG19. There were two RDMs—the RDM of the layer conv5_4 (the last convolutional layer) and the RDM of the layer fc2 (the last fully connected layer before SoftMax layer). As the RDM of the layer conv4_2, the two candidate RDMs were constructed using the corresponding feature maps extracted from the VGG19 by the 50 images from the testing set. We selected the correlation distance (1 minus the linear correlation between each pair of feature maps) as the measurement of representational dissimilarity to construct each RDM.

The second type were RDMs derived from brain activity. We used the measured brain activity from the testing sets and the imagery sets to constructed RDMs, respectively. Because the categories of the stimuli (nouns) in the imagery data were the same as the categories of the stimuli (images) in the test data, the RDMs from the imagery data could be treated as RDMs using the same set of stimuli—the 50 images from the test data. There were 14 RDMs for each subject, 7 RDMs from the test data for each ROI and 7 RDMs from the imagery data for each ROI (V1, V2, V3, V4, LOC, FFA, and PPA).

The third type were RDMs derived from stimuli (50 images from the testing set) directly. There were three RDMs—the RDM of Gabor features, the RDM of silhoutee, and the RDM of semantics. The RDM of Gabor features was constructed using the Gabor features of images extracted from the GWP model. To construct the RDM of silhoutte, we converted images to silhouettes (binary images in which each figure pixel is 0 and each background pixel is 1) and calculated the correlation distance between each pair of silhouettes. To construct the RDM of semantics, we calculated the semantic distance between each pair of images. We used the function “path_similarity” from the Natural Language Toolkit library [53] to calculate how similar two categories of images are (semantic similarity), which based on the WordNet [54]. The score returned from the function was in range 0 to 1, so we converted the semantic similarity to the semantic distance by subtracting the score from 1.

In total 19 candidate RDMs were constructed to compare with the reference RDM for each subject and the Spearman’s rank correlation coefficient was selected to measure the similarity between each candidate RDM and the reference RDM. After that, we performed statistical inference to answer two questions—whether a candidate RDM and the reference RDM were significantly correlated (by permutation test with 10000 permutations) and whether the correlation between a candidate RDM and the reference RDM was significantly different from the correlation between another candidate RDM and the reference RDM (by bootstrap test with 1000 replications). For each test, FDR was applied for multiple comparison correction [49]. All calculations were done using MATLAB 2020a and Python 3.7 on a Linux (Ubuntu 18.04 LTS) desktop with a Geforce GTX 1660 Ti graphics card (6 Gb of VRAM).

## 3. Results

### 3.1. The deep features which represented the semantic content of an image mainly predicted the activity of voxels in the early visual areas

Because there were no models survived in the imagery set, the following analysis mainly focused on the testing set. The number of survived models (voxels) in the testing set was 297 of 4466 for S1, 539 of 4404 for S2, 1165 of 4643 for S3, 1041 of 4133 for S4, and 561 of 4370 for S5. The distributions of survived models and corresponding prediction accuracy between ROIs were different among subjects (Fig.2.A and for Fig.2.B for S1, the result of other subjects can be found in Appendix, same below in all figures). But we still observed some clear common trends: the deep features extracted from the conv4_2 layer of the VGG19, which were selected as the semantic content representation of an image in the original NST algorithm, mainly predicted the activity of voxels in the early visual areas (V1, V2, and V3). First, most of the survived voxels located in the early visual areas for each subject (99% for S1, 90% for S2, 78% for S3, 80% for S4, and 91% for S5); Second, the prediction accuracy for early visual areas were higher than that of higher visual areas.

**Figure 2:**
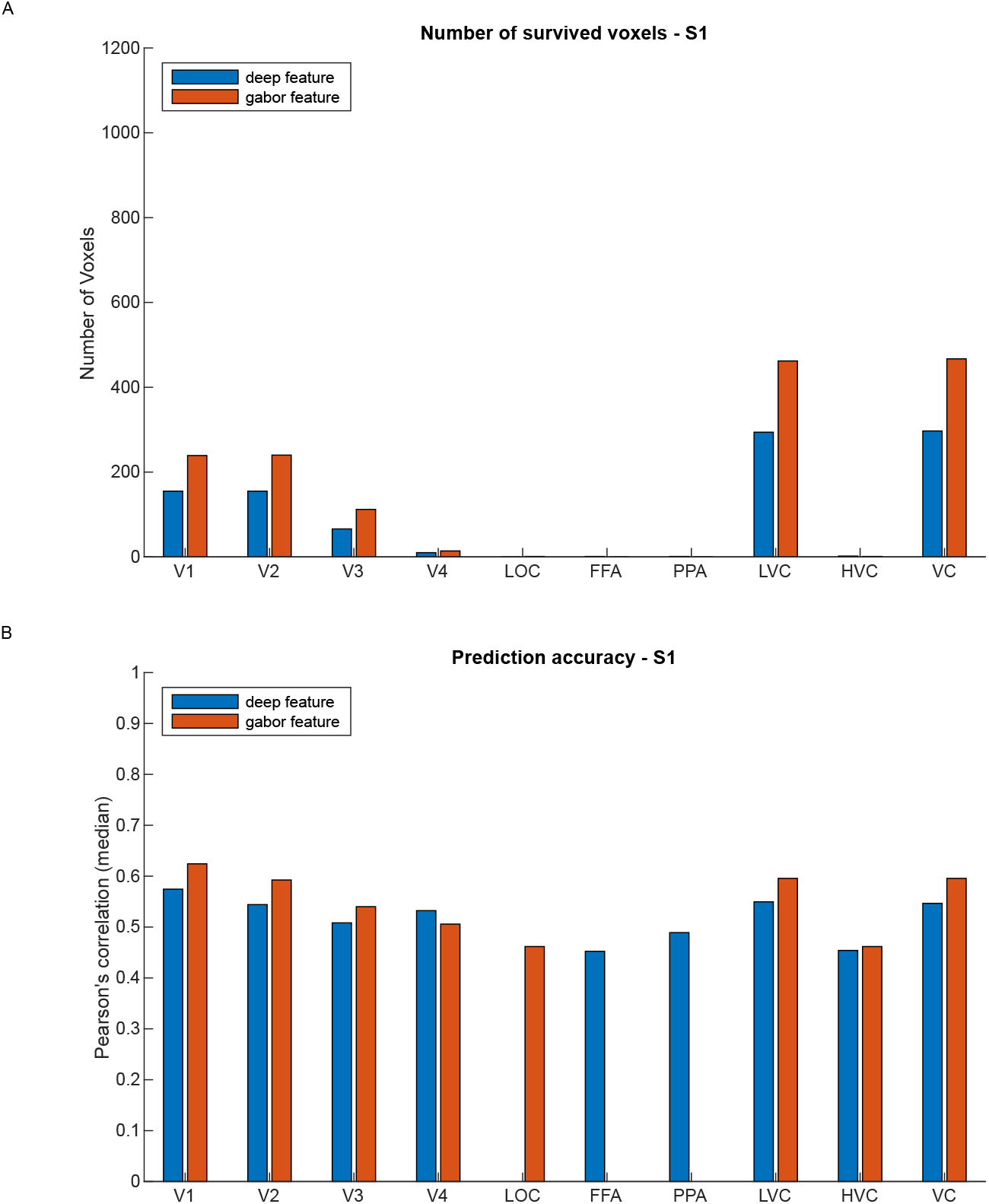
The result of encoding models based on deep features and gabor features for S1. (A) The number of survived voxels for each ROI. (B) The prediction accuracy for each ROI (the median of Pearson’s correlation coefficients of all survived voxels in each ROI).

The survived models could be used to decode stimuli from the measured brain activity—image identification using the testing set (Fig.3.A for S1). The identification accuracies of 5 subjects were 94% (47/50), 88% (44/50), 100%, 100%, and 92% (46/50). After checked all the identification errors, we founded that there were some common mistakes among different subjects. All the 4 images (No.17, No.19, No.41, and No.44) that were incorrectly identified by the encoding models of S5 were also incorrectly identified in S2, and three of them (No.19, No.41 and No.44) were incorrectly identified in S1 too. The encoding models of S1 made the same mistake as the models of S2, which identified the No.41 image as the No.42 image. And the encoding models of S2 made the same mistake as the models of S5, which identified the No.44 image as the No.26 image and the No.17 image as the No.22 image (For copyright reasons, we can not show the actual images).

**Figure 3:**
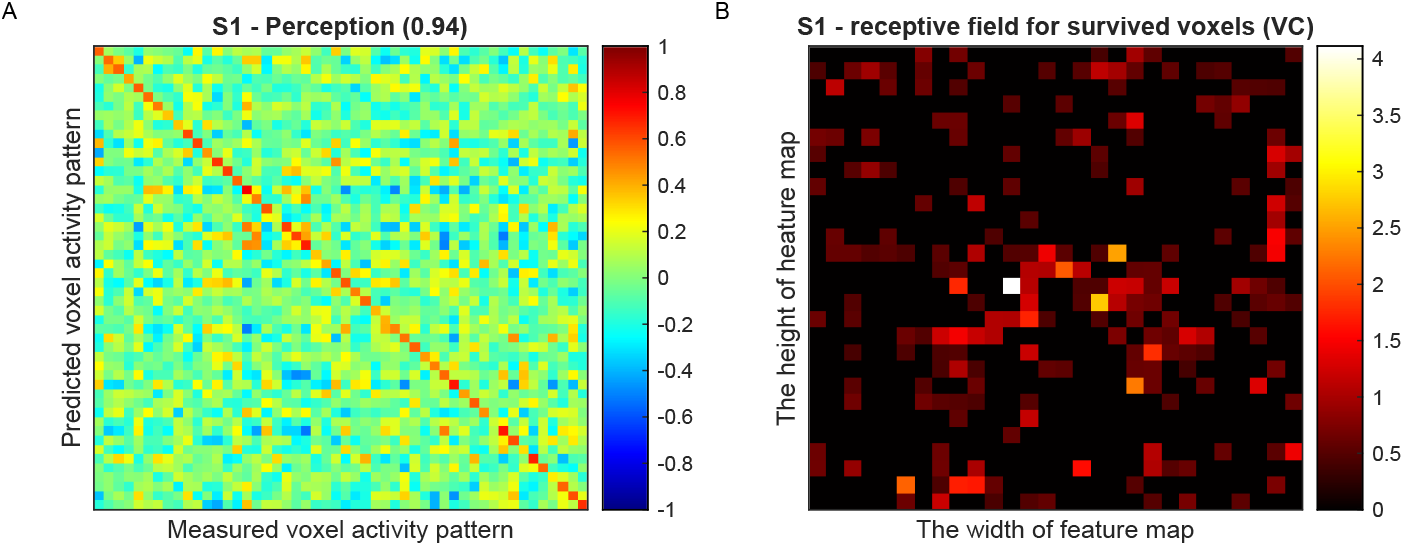
Decoding performance and receptive field analysis based on encoding models using deep features for S1. (A) The decoding performance for S1 (identifying stimuli from measured brain activity using the testing set). (B) The distribution of receptive fields for survived voxels on the feature map. The value of each location equaled the sum of prediction accuracy (r) for all survived voxels located in that location.

Because Lasso regresssion enables feature selection, the survived models also described the relationship between features (*X*) and voxel responses (*y*) through regression coefficients. From the perspective of voxels, we calculated the number of features each ROI related (median) and found no common trend among subjects. From the perspective of features, we calculated the number of voxels each feature related and ranked features according to the number of voxels its related. Then we calculated Pearson’s correlation coefficients of ranked feature index between each subject pair to examine if there were similar patterns among subjects. The result showed that there was no significant correlation. But if only considered the top-5 features, we found that most of the features were same among subjects (Table1).

**Table 1:** Top-5 features (Index)

| Feature Order | Subject1 | Subject2 | Subject3 | Subject4 | Subject5 |
| --- | --- | --- | --- | --- | --- |
| 1 | 383 | 250 | 250 | 383 | 383 |
| 2 | 60 | 60 | 383 | 60 | 250 |
| 3 | 250 | 289 | 60 | 250 | 60 |
| 4 | 289 | 383 | 355 | 448 | 482 |
| 5 | 355 | 198 | 491 | 289 | 289 |

The features (*X*) of each survived model corresponded to a spatial location on the feature map (we only used the features from one spatial location of the feature map as *X*), which could be seen as the center of population receptive field of the related voxel (*y*). So we could visualize the distribution of receptive fields of survived voxels for each subject (Fig.3.B for S1). The result showed that survived voxels distributed widely on the feature map and there was a slight trend that some of voxels clustered near the center of the feature map.

### 3.2. The decoding performance was better when encoding models used the deep features

Previous study showed that Gabor features also predicted the activity of voxels in the early visual areas [50]. So we constructed encoding models based on Gabor features and compared it with encoding models using the deep features. Like the encoding models using the deep features, no voxels survived in the imagery set when encoding models used gabor features. So we only compared two different types of encoding models using the test set. From the perspective of individual voxel, there were more voxels survived in the early visual areas with encoding models based on Gabor features for all subjects(Fig.2.A for S1). And the prediction accuracy of the early visual areas (the median of Pearson’s correlation coefficients of all survived voxels in each ROI) was also higher for all subjects when encoding models used Gabor features (Fig.2.B for S1). It suggested that, relative to the deep features, Gabor features were preferentially represented by the early visual areas. From the perspective of activity pattern of voxels (decoding performance), however, the decoding performances were better for all subjects when encoding models used the deep features (Fig.4.A for S1).

**Figure 4:**
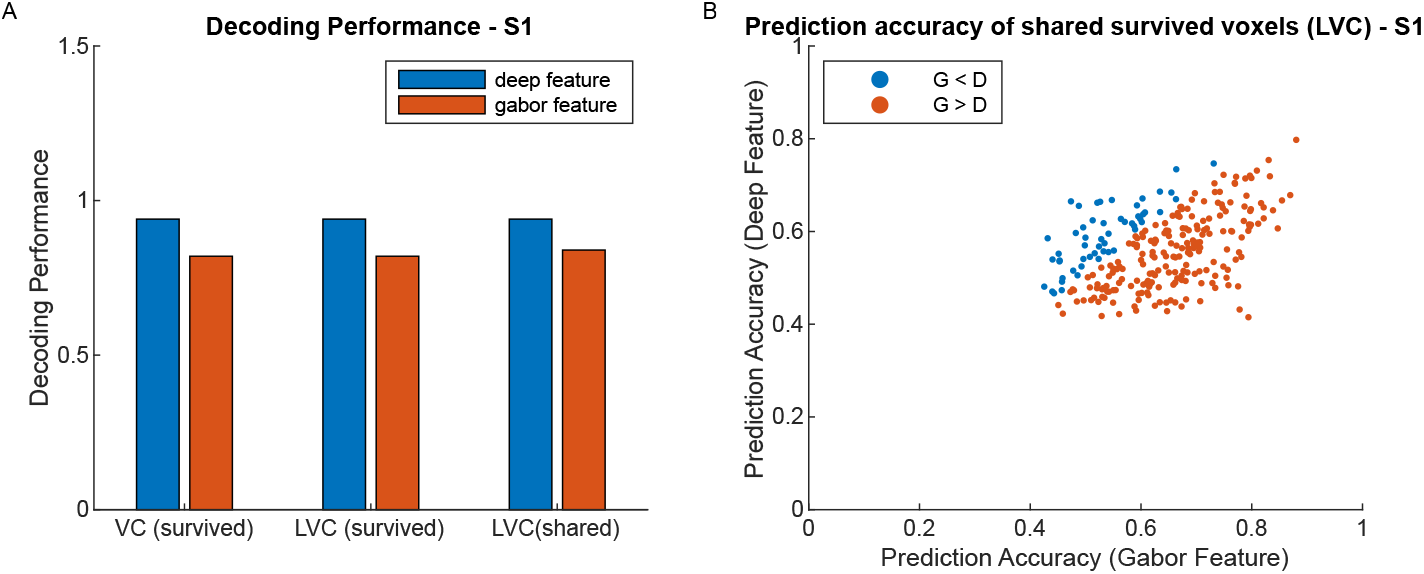
The comparison of two types of encoding models for S1. (A) Comparison of decoding performance for S1. VC (survived) = all survived voxels, LVC (survived) = all survived voxels in the early visual areas, LVC (shared) = voxels simultaneously survived in the early visual areas for both models. (B) The prediction accuracy of shared voxels (voxels simultaneously survived in the early visual areas for both models) for S1.

The better decoding performance of encoding models using the deep features could be due to the survived voxels in the higher visual areas (there were more survived voxels in higher visual areas when encoding models used deep features), so we excluded survived voxels not in the early visual areas for both models and compared identification performance again. The results showed that, for all subjects, the identification performances were still better when encoding models used the deep features (Fig.4.A for S1). Then we chosed voxels simultaneously survived in the early visual areas for both models and found that the result did not change—the identification performances were better when encoding models used the deep features for all subjects (Fig.4.A for S1). Further analyses on these voxels, we observed a common trend among all subjects—the prediction accuracies of shared voxels showed a positive correlation between two types of models and more voxels were better predicted when encoding models used Gabor features (Fig. 4.B for S1).

### 3.3. The representation of the semantic content of an image did related to the semantics of the image and also preserved visual details of the image to some extent

In accordance with the analysis using encoding models, the RSA was also based on individuals (Fig.5 for S1). The RDMs from the pre-trained VGG19 were significantly correlated with the RDM of the layer conv4_2. For all subjects, the RDM of the layer conv5_4 (the last convolutional layer) showed the strongest correlation (*r_s_* = 0.49) and the RDM of the layer fc2 (the last fully connected layer before SoftMax layer) was in the second echelon among all candidate RDMs (*r_s_* = 0.31). This was a reasonable result given that all three RDMs derived from the same model (VGG19) and the layer conv5_4 was more similar to the layer conv4_2 than the layer fc2.

**Figure 5:**
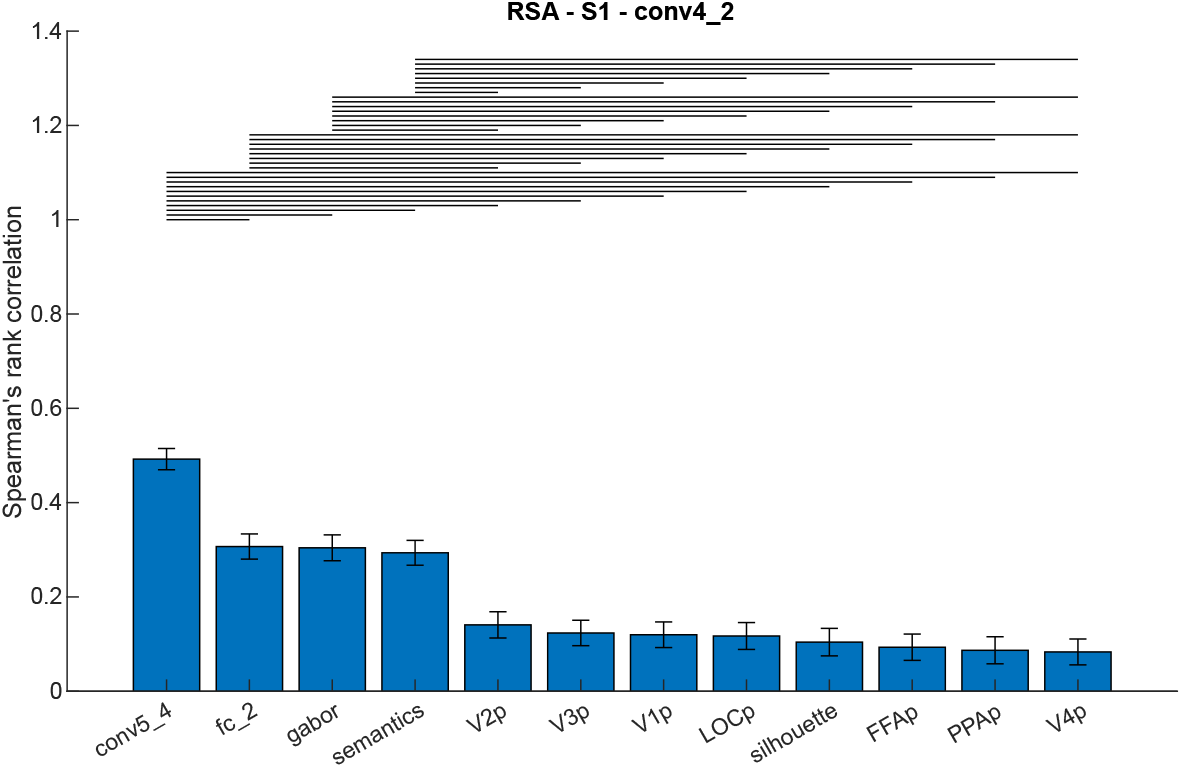
The result of RSA for S1. A horizontal line over two bars indicates that the difference of correlation is significant.

The RDMs from the stimuli were significantly correlated with the RDM of the layer conv4_2, too. For all subjects, the RDM of Gabor features (*r_s_* = 0.30) and the RDM of semantics (*r_s_* = 0.29) were both in the position of the second echelon. It suggested that the representation of the layer conv4_2 did relate to the semantics of stimuli and also preserved visual details of stimuli to some extent. The RDM of silhouette was in the position of the lowest echelon for all subjects (*r_s_* = 0.10). Because the silhouette of an object provided a limited description of the specific shape of the object, the representation of the layer conv4_2 also related to the specific shape of stimuli.

The situation of the RDMs from brain activity was more complex. For each subject, all 7 RDMs from the testing sets were significantly correlated with the RDM of the layer conv4_2. Although there were some individual differences about the relative position of these RDMs, the RDMs from the early visual areas which were in the second or third echelon roughly showed stronger correlation than those from the higher visual areas. In contrast, there were few RDMs from the imagery sets significantly related to the RDM of the layer conv4_2 (LOC for S5, which was in the position of the lowest echelon). This result was in line with the result of encoding models to some extent.

## 4. Discussion

The impenetrability of DNNs reduced the explanatory power of studies which used DNNs as computational models to explore the biological brain. Inspired by NST, we circumvented this problem by using deep features that were given clear meaning—the representation of the semantic content of an image. Using encoding models, we quantitatively showed that the deep features which represented the semantic content of an image mainly predicted the activity of voxels in the early visual areas. It was a surprise that the semantics-related features mainly predicted the voxels in the early visual areas rather than those in the higher visual areas. Then we compared encoding models using the deep features with encoding models using Gabor features which have been proved to also predict the activity of voxels in the early visual areas [50] and found that the decoding performance was better when encoding models used the deep features, which suggested that the deep features might contain more information about stimuli than Gabor features. These results naturally led us to another question: what the representation of the semantic content of an image really is?

The results of RSA showed that, the representation of the semantic content of an image did related to the semantics of the image and also preserved visual details of the image to some extent, which suggested that the representation of the semantic content of an image might be a hybrid form—both in propositional format and depictive format. It was a reasonable result from the perspective of NST—although there is no clear definition of the semantic content of an image, it is easily understood that to make NST successful, the representation of the semantic content of an image should preserve visual details of the object in the image to some extent, which guaranteed the generated images described the same thing.

But how could the format of a representation be both propositional and depictive? The question of how the information is represented in our brain had been discussed for many years, which was known as the imagery debate [55]. At the heart of the debate was whether all information is represented in a symbolic, propositional format. Convergent evidence from empirical studies of mental imagery suggested that information can be represented in a pictorial, depictive format [56]. The existence of the depictve format of information ended the imagery debate but also raised new questions: how many formats can the brain use and what is the relationship between these formats and the propositional format? With the development of theories of grounded cognition, the dominant position of the propositional format in cognition is being challenged. From the perspective of grounded cognition, there were no amodal symbols in our brain that were independent of the modal representation and all cognitive phenomena were ultimately grounded in modal simulations, bodily states, and situated action, which was supported by many researches on perception, memory, language, thought, social cognition, and development [57–59]. This view emphasized the key role of the depictive, modality-specific representation in cognition and denied the independent existence of the symbolic, propositional representation, which was clearly articulated by Comenius from several hundred years ago—“things are essential, words only accidental; things are the body, words but the garment; things are the kernel, words the shell and husk. Both should be presented to the intellect at the same time, but particularly the things, since they are as much objects of understanding as is language” [60, p. 435].

From this view, the representation of information in our brain is essentially depictive and can implement symbolic functions naturally. This is in line with our result to some extent. On the one hand, the representation of the semantic content of an image (the feature maps of the layer conv4_2) was essentially depictive. This was because the feature maps extracted from the convolutional layer of the VGG19 naturally preserved the topology of the original image. Besides, The reuslt of RSA also showed that it preserved visual details of the image. On the other hand, this representation did reflect the semantics of the image in some degree.

In fact, there was another theory also addressing the relationship between the propositional representation and the depictive representation of information in our brain—the dual coding theory [61], which emphasized the beneficial effects of the depictive representation of information on cognition (concreteness) and suggested that the two types of representations are independent from each other in our brain (Paivio believed that there were two distinct subsystems in our brain specialized for dealing with different types of representations). Both theories admitted the association between the two types of representations but disagreed with each other about the relationship between the two types of representations. From our result, we tend to support the monistic view of the two types of representations. But we also noticed that our result only involved the early visual areas which were is the early stages of the visual ventral stream. So How does the depictive representation of an object change along the visual ventral stream to make object recognition possible—whether the independent propositional representation will eventually appear—is not clear. This question needs further studies.

Unlike the previous study [14], we did not observe that encoding models which were trained using the perceptual data could successfully predict voxel responses from the imagery data. This could be due to different experimental tasks. In the study of [14], subjects were asked to imagine particular artworks, such as “Betty” by Gerhard Richter and “Horse Bath” by Odd Nerrdum. In the study of [16], which provided data for this paper, subjects were asked to imagine as many object images as possible from concrete nouns, such as leopard and swan. The difference between two tasks was whether the imagery had a particular content. For example, when you were asked to draw your cat or dog, what you drew must be a particular cat or dog; but when you were asked to draw a cat or dog, you could draw any cat or dog, even Hello Kitty or Snoopy. Because of the individualization and arbitrariness of the imagery in our study, it seems reasonable that our result was not consistent with the previous study and could not address the issue of the relationship between perception and imagery.

In addition to the obvious individual divergences in encoding mechanisms, our result showed that there were some common mechanisms among subjects (common mistakes in image identification and similarity of the top 5 features). In contrast to the symbolic, propositional representations, the depictive, modality-specific representations of information were grounded in the modalities, the body, and the environment. So they were highly personal and changed from time to time. This was a key difference between the grounded cognition and traditional cognitive theories and could be used to explain individual divergences in cognition. Meanwhile, we did share a common physical and environmental basis, which was also reflected in cognitive process and made communication possible. This may explain the existence of the common mechanisms.

## 5. Conclusions

In this study, we quantitatively showed that the deep features which were selected as the representation the semantic content of an image in the original NST algorithm mainly predicted the activity of voxels in the early visual areas and these features were essentially depictive but also propositional. This results implied that some depictive representation of an object in our brain can naturally reflect semantics of the object to some extent and this phenomena can be found in the early visual areas, which provided empirical evidence to the core viewpoint of the ground cognition.

## Acknowledgments

This work was supported by the National Natural Science Foundation of China (31600907).

## Author contributions

Ming Liu, Delong Zhang, and Huawei Xu conceived the study; Delong Zhang and Huawei Xu performed the analysis; Delong Zhang and Huawei Xu wrote the manuscript. All authors participated in discussion and approved the final version of the manuscript.

## Appendix. Supplementary materials

Supplementary materials to this article can be found online at

## Supplementary Materials

**Fig.S1.**
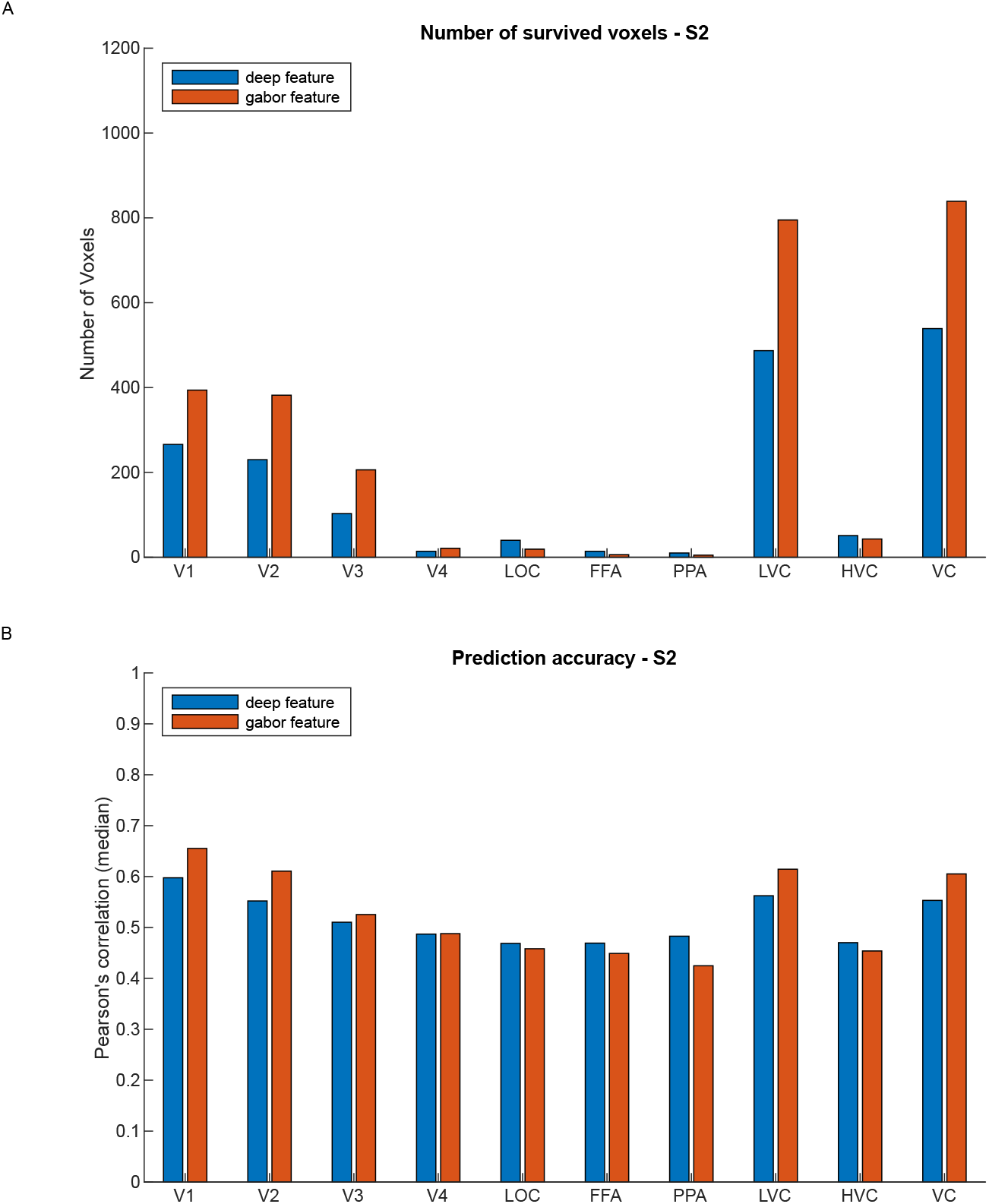
The result of encoding models based on deep features and gabor features for S2.

**Fig.S2.**
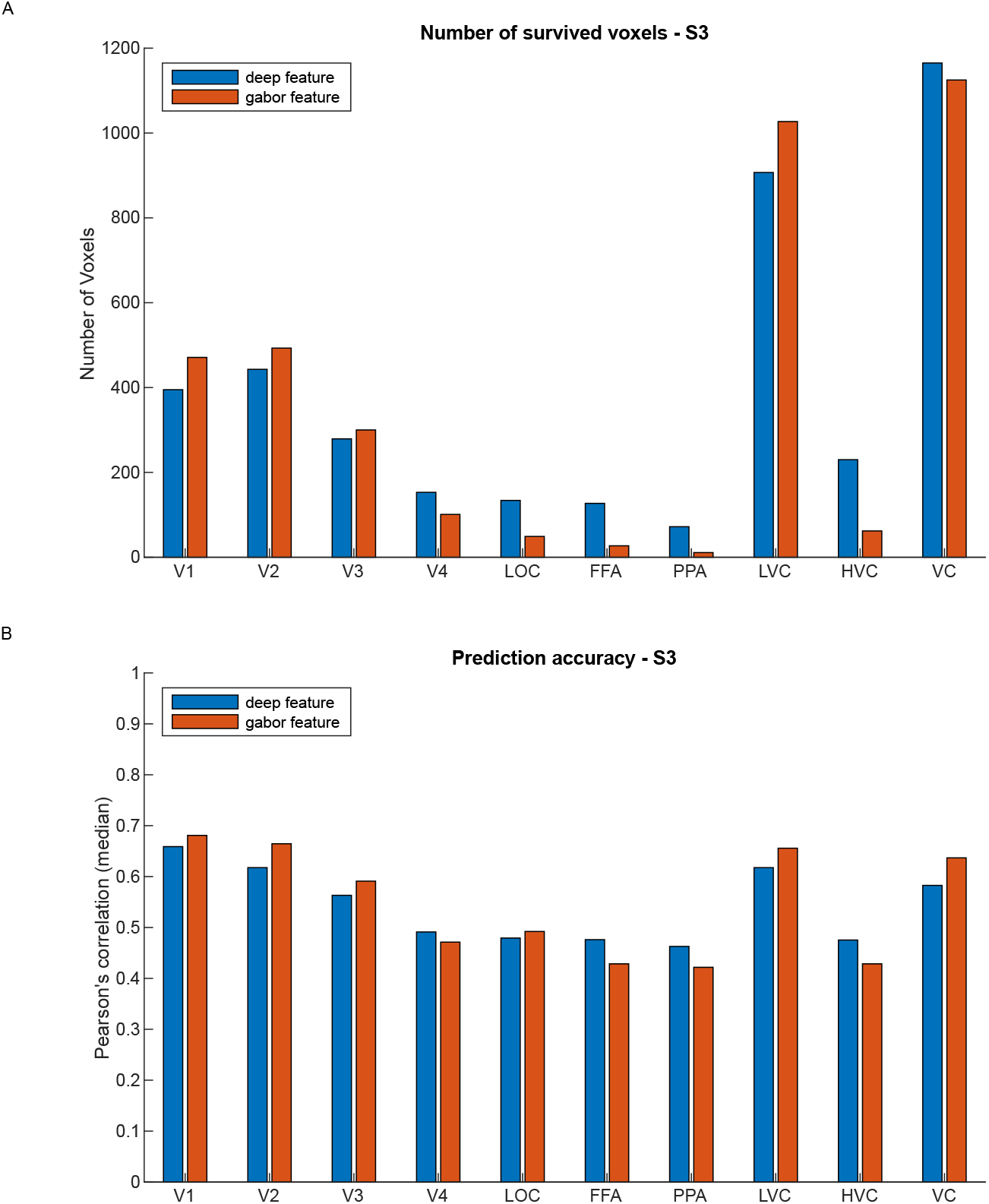
The result of encoding models based on deep features and gabor features for S3.

**Fig.S3.**
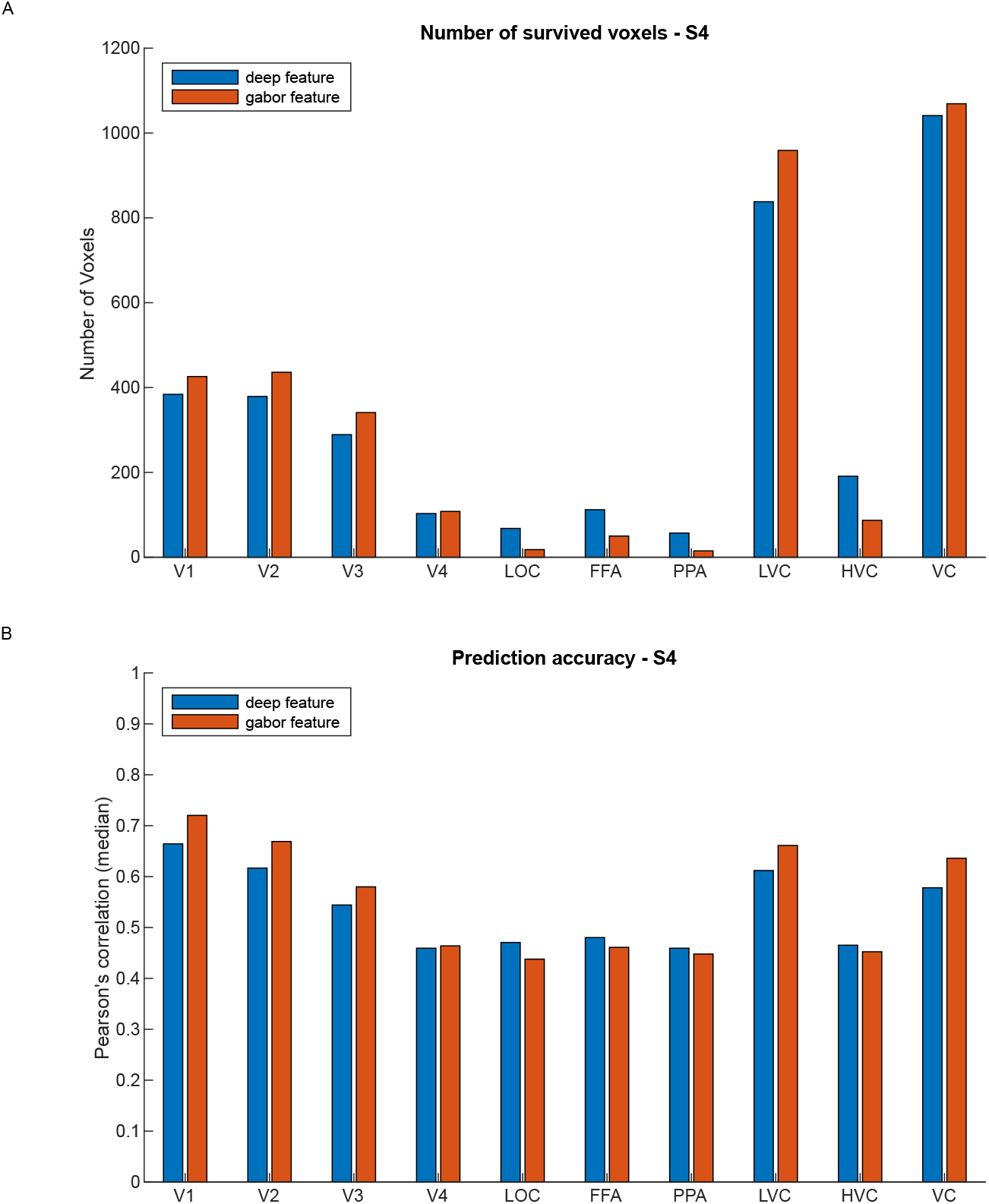
The result of encoding models based on deep features and gabor features for S4.

**Fig.S4.**
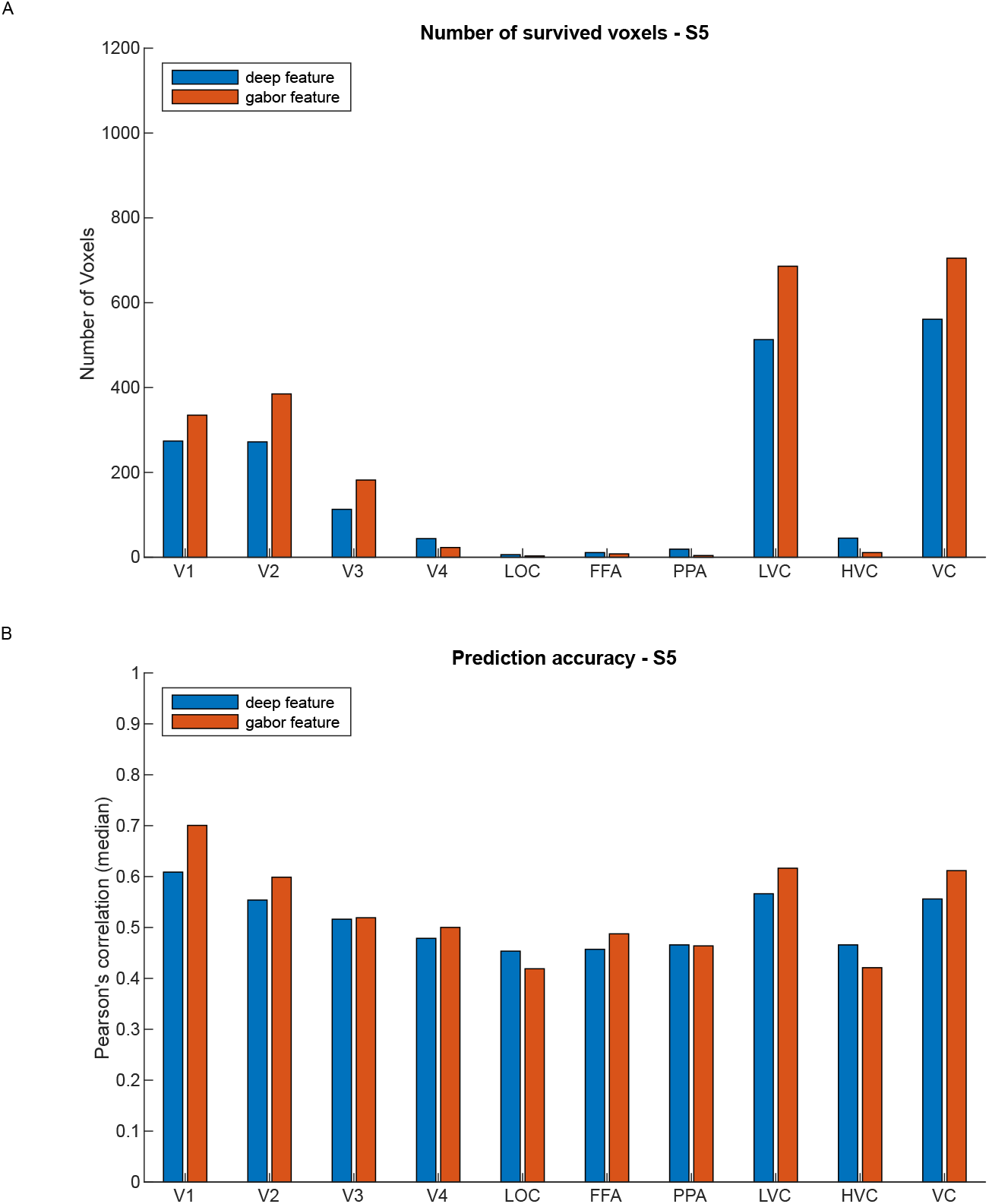
The result of encoding models based on deep features and gabor features for S5.

**Fig.S5.**
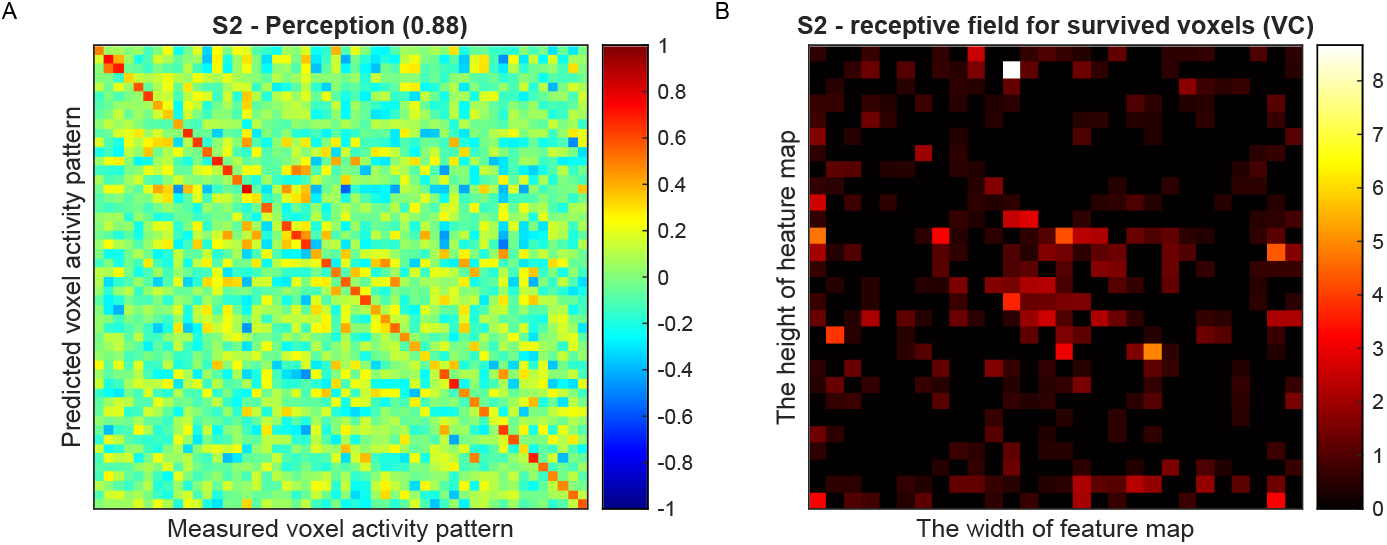
Decoding performance and receptive field analysis based on encoding models using deep features for S2.

**Fig.S6.**
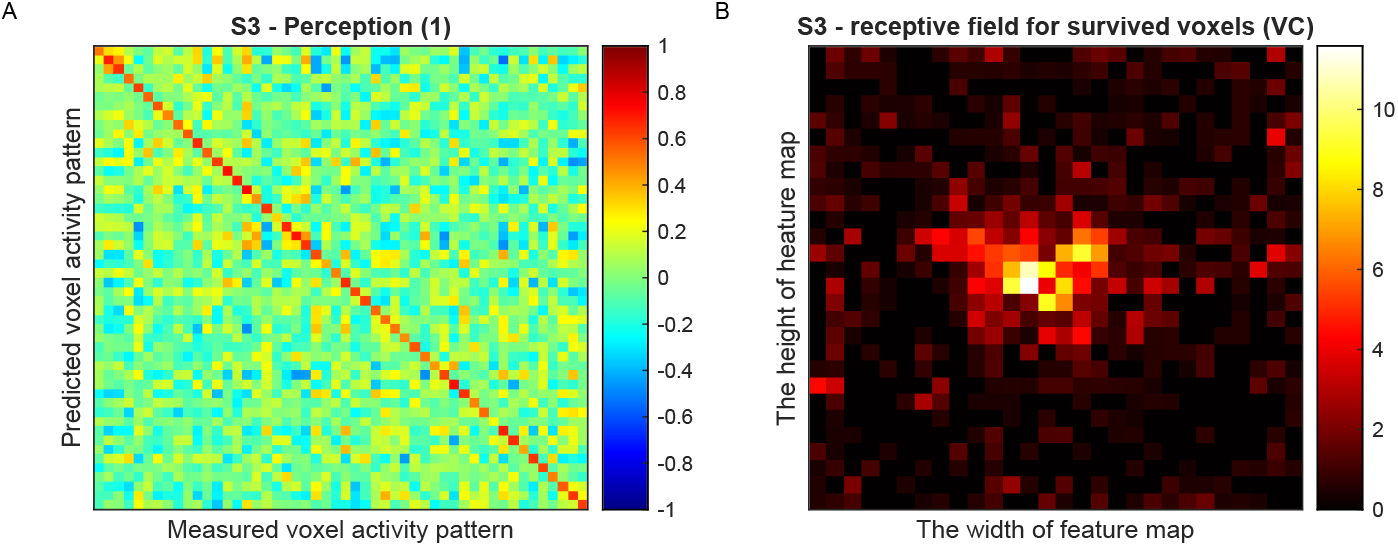
Decoding performance and receptive field analysis based on encoding models using deep features for S3.

**Fig.S7.**
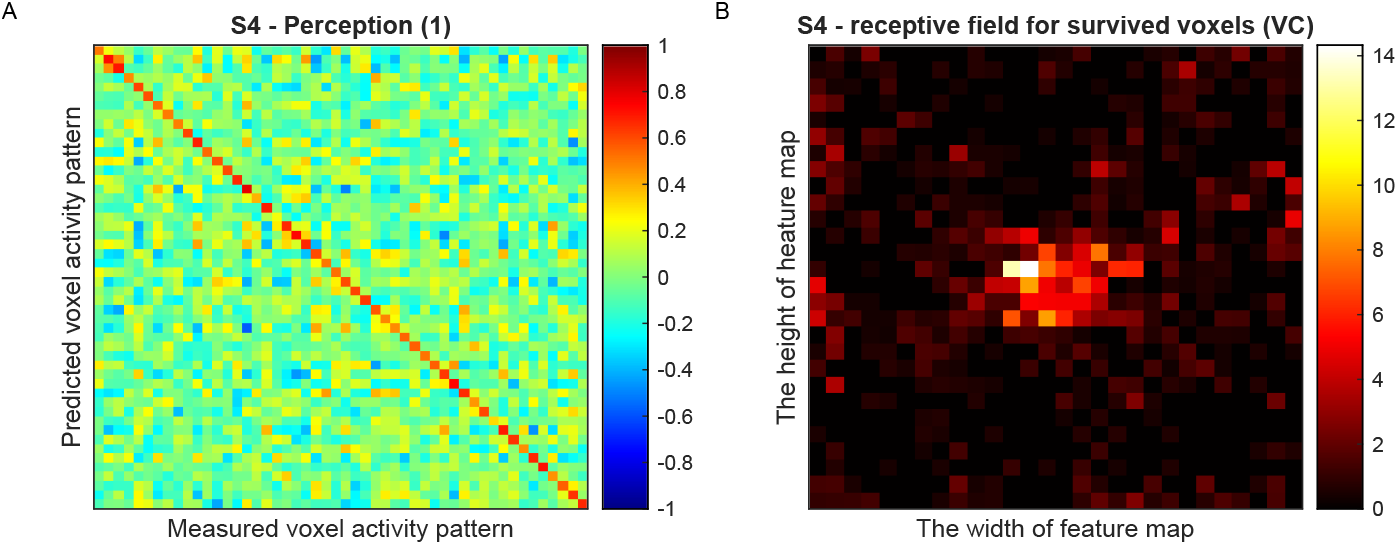
Decoding performance and receptive field analysis based on encoding models using deep features for S4.

**Fig.S8.**
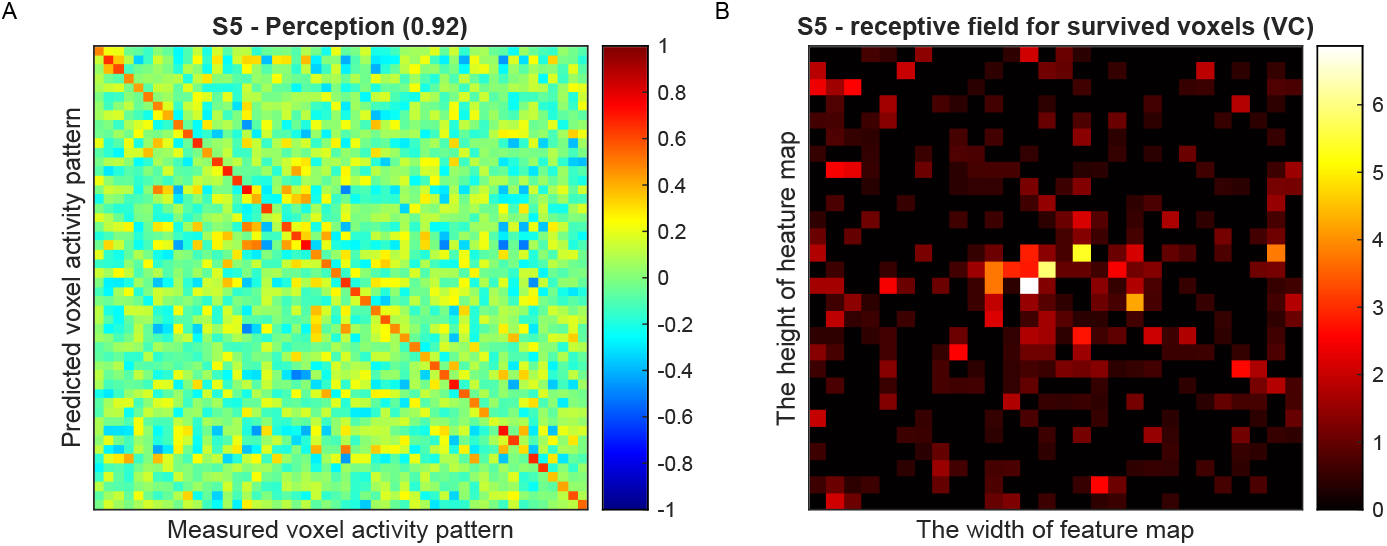
Decoding performance and receptive field analysis based on encoding models using deep features for S5.

**Fig.S9.**
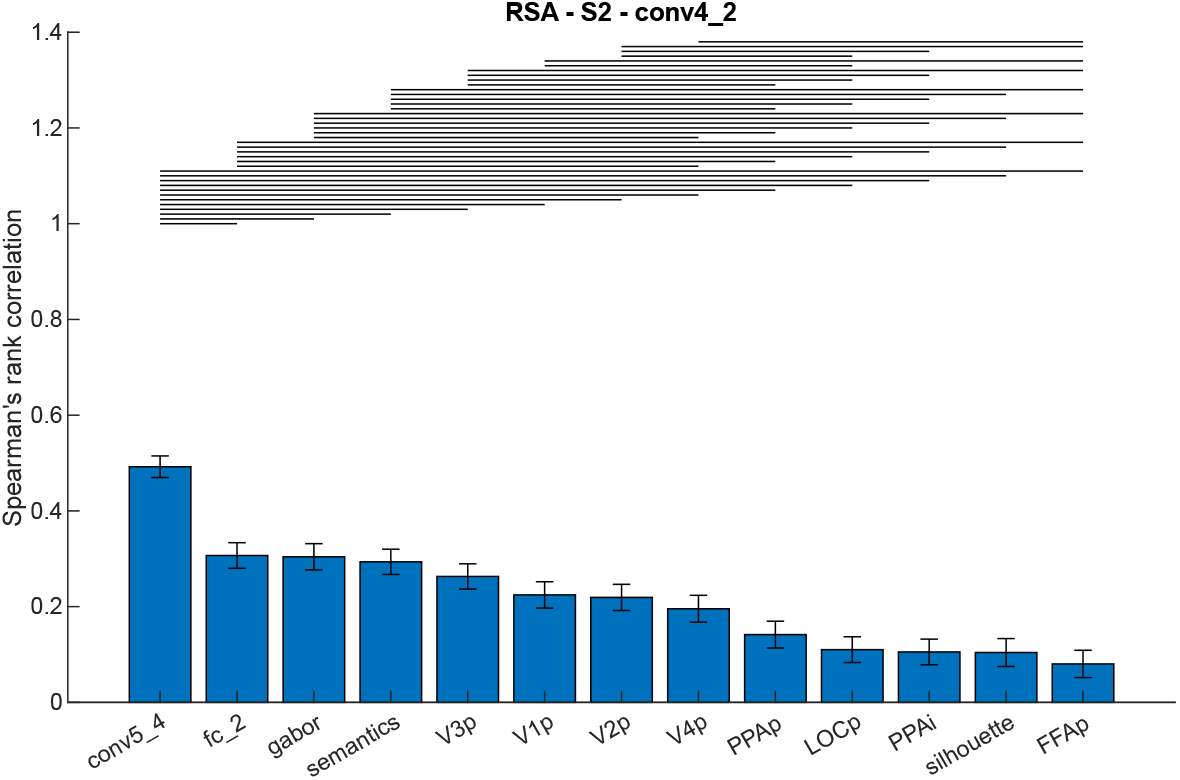
The result of RSA for S2.

**Fig.S10.**
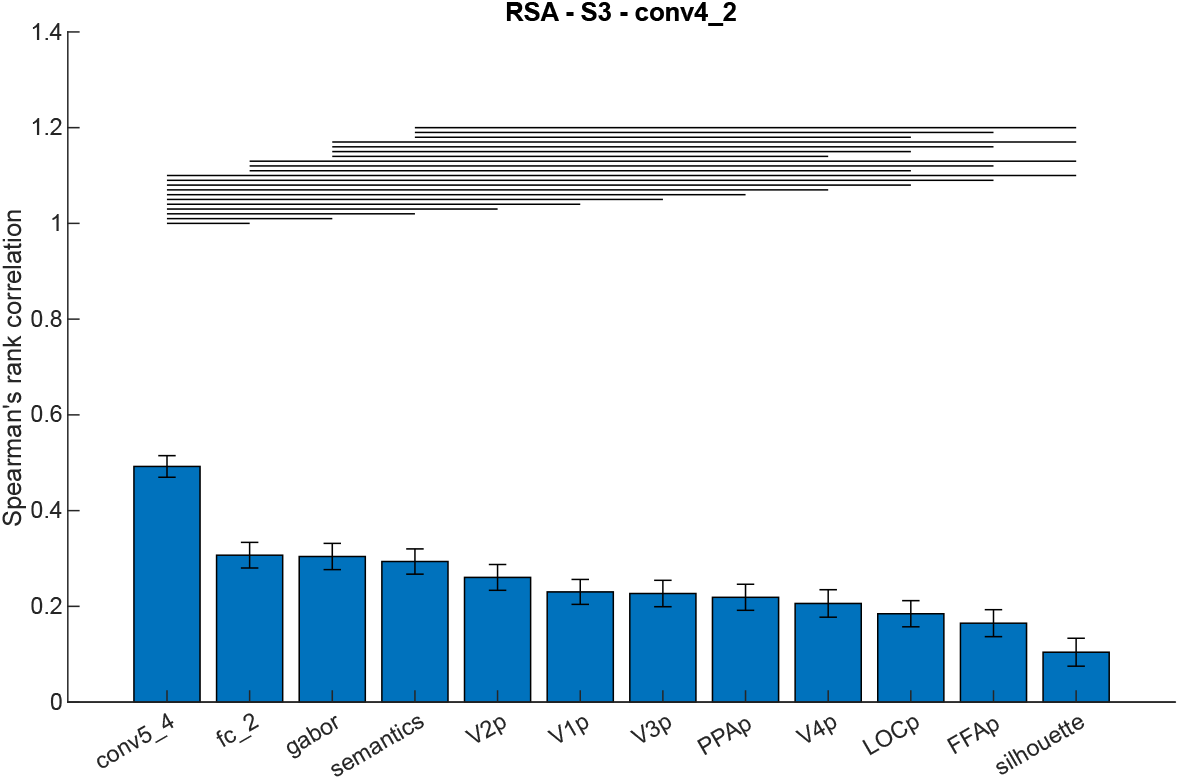
The result of RSA for S3.

**Fig.S11.**
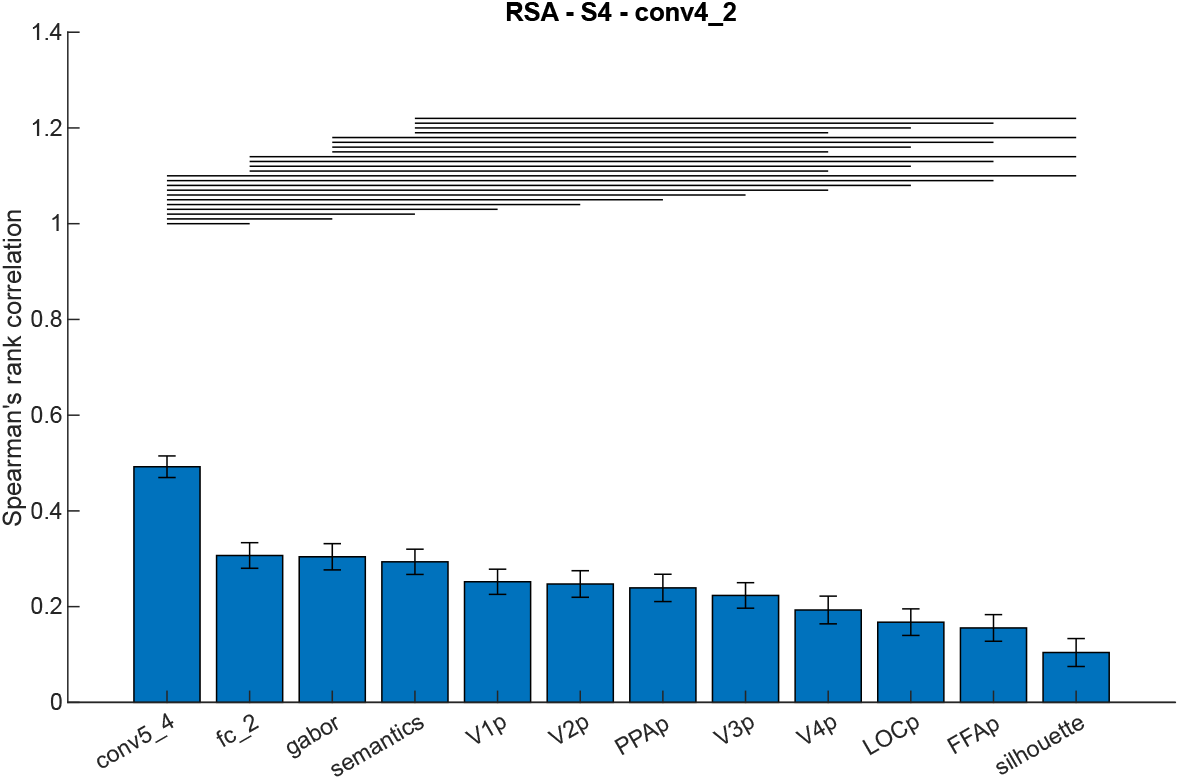
The result of RSA for S4.

**Fig.S12.**
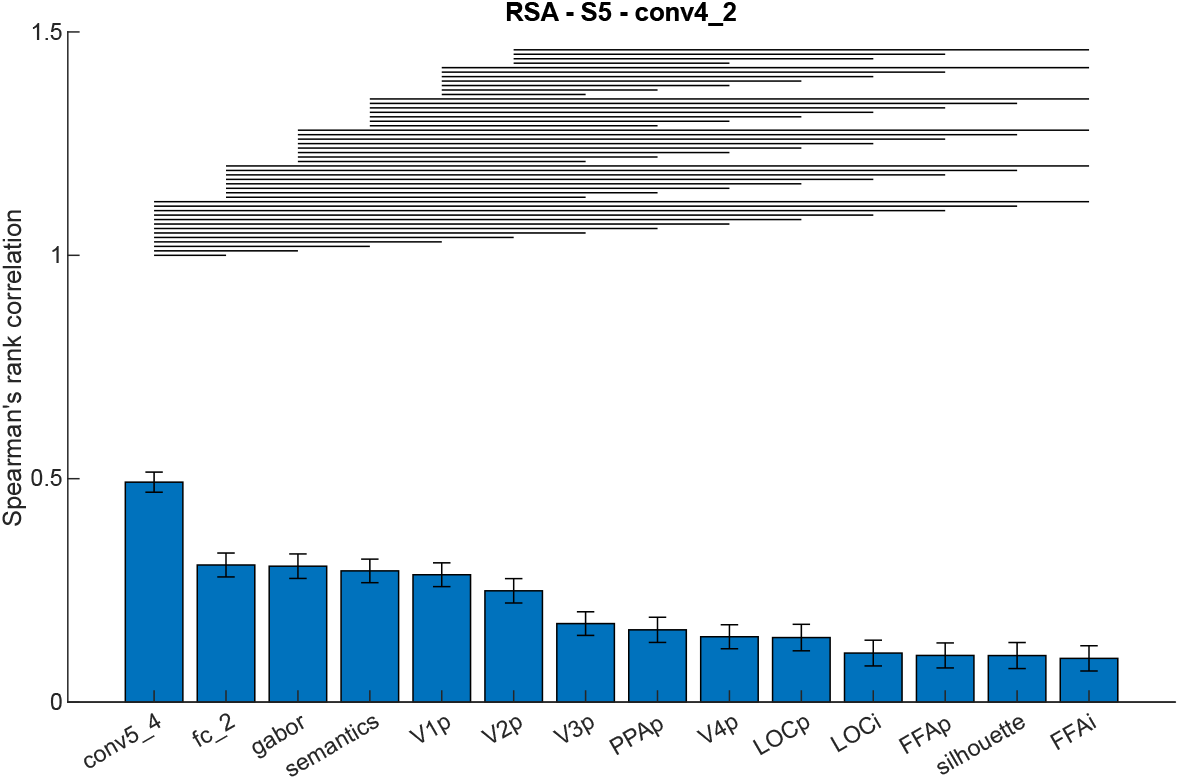
The result of RSA for S5.

